# Susceptibility of Glucose Regulation to Social Isolation

**DOI:** 10.64898/2025.12.18.695168

**Authors:** Hannah H. Lamont, Rumi Oyama, Paula Diaz Munoz, Dashiel Siegel, Paula Baringanire, Valentina Vargas, Miriam E. Bocarsly, Vanessa H. Routh, Ioana Carcea

**Author notes:** equal contribution (Hannah H. Lamont, Rumi Oyama).

## Abstract

Loneliness and social isolation strongly associate with increased incidence of diabetes in humans. However, it remains unclear if lack of meaningful social interactions represents a cause or a symptom of disease. In rodents, social isolation leads to metabolic dysregulation, however the dynamics and contributing factors remain poorly understood. Here we show that single-housing young adult male mice for at least three weeks led to fasting hyperglycemia, an effect that was maintained for the duration of single-housing, but was corrected within a week of reverting mice to co-housing conditions. Single-housing did not affect glucose regulation in females. Gonadectomy experiments revealed that testicular factors induced susceptibility to social isolation, as orchiectomy prevented isolation-induced fasting hyperglycemia in males. We did not find a protective role for ovarian hormones, as ovariectomized females were as resilient as intact females to isolation-induced fasting hyperglycemia. To understand the underlying mechanisms for susceptibility to isolation, we measured plasma levels of glucoregulatory hormones. Isolation did not affect the levels of insulin, epinephrine, and corticosterone. However, glucagon levels were distinctly modulated by social isolation in intact and orchiectomized mice. Social isolation induces increased immediate early gene expression in specific neurons of the ventromedial hypothalamus, a glucoregulatory brain structure that promotes glucagon release. Taken together, our findings show that testicular hormones make males susceptible to isolation-induced disruption of glucose regulation and suggest that brain glucoregulatory neurons play a role.

## Introduction

Longitudinal and meta-analytical studies in humans identified a strong link between social contexts and health. Supportive social environments associate with increased longevity and better health status, whereas isolation and perceived loneliness associate with higher mortality rates and increased incidence of mental disorders, metabolic and cardiovascular disease, and immune dysfunction (Chen et al., 2024; Holt-Lunstad et al., 2015; Holt-Lunstad & Steptoe, 2022; Office of the Surgeon, 2023; Song et al., 2023). However, it remains unclear if the relationship between social environments and health is causal, as several studies analyzing observational human data reported contradictory findings (Liang et al., 2024; Song et al., 2023).

Rodent animal models have been used to determine if manipulating social environments affects physiological measures of wellbeing. Social isolation affects several metabolic endpoints in mice and rats: weight, body composition, glucose, and glucoregulatory hormones (Bove et al., 2022; Chen et al., 2025; Mountain et al., 2023; Nonogaki et al., 2007; Smolensky et al., 2024; Sun et al., 2014). We recently found that isolated male mice were refractory to insulin-induced hypoglycemia (Patel et al., 2023). However, both an in-depth characterization and a mechanistic understanding of how social isolation affects glucose regulation are lacking. Moreover, although sex differences in central glucose sensing exist (Santiago et al., 2016a, 2016b), studies of potential sex differences in socially-regulated glucose levels are lacking.

Blood glucose levels are tightly regulated by multiple organs that coordinate their function to maintain glucose homeostasis via hormonal and autonomic processes. Major roles are played by glucose sensing cells in the pancreas and the brain. Pancreatic α-cells secrete the gluconeogenic and glycogenolytic hormone glucagon in response to decreasing glucose levels, whereas ß-cells secrete insulin in response to increasing glucose levels (Frohman, 1969; Thorens, 2022). Adrenal hormones epinephrine and glucocorticoids, and the autonomic system also play important roles in glucose regulation (Hoffman, 2007; Vinson, 2009).

A network of glucose-responsive brain structures modulate autonomic input to the pancreas, regulating the release of insulin and glucagon (Rosario et al., 2016; Thorens, 2022). Some of these structures have been implicated in social behavior as well and represent potential integrators of social and glucoregulatory functions. For example, oxytocin neurons in the paraventricular nucleus of the hypothalamus (PVN) regulate pancreatic hormonal activity (Papazoglou et al., 2022), and have been implicated in many social behaviors (Carcea et al., 2021; Ferretti et al., 2019; Froemke & Young, 2021). Moreover, acute social isolation increases the proportion of oxytocin neurons in PVN as well as immediate early gene expression in these neurons (Musardo et al., 2022). The PVN is anatomically and functionally connected with the ventrolateral division of the ventromedial hypothalamus (vlVMH), another brain structure with key roles in both glucose regulation and social behavior (Borg et al., 1997; Borg et al., 1995; Falkner et al., 2016). vlVMH glucose-inhibited and glucose-excited neurons regulate sympathetic to parasympathetic balance, and can send polysynaptic inputs to the pancreas (Hirschberg et al., 2020; Rosario et al., 2016; Routh, 2010; Shimazu & Minokoshi, 2017). Notably, vlVMH neurons that express neuronal nitric oxide synthase (nNOS) are activated by decreased glucose (Fioramonti et al., 2010; Fioramonti et al., 2011), and their activation increases circulating glucagon and blood glucose (Faber et al., 2018). It remains unclear if and how these glucose-regulatory mechanisms are affected during chronic social isolation.

In our study, we show that social isolation induces progressive but reversible fasting hyperglycemia in male but not female mice. We find that male gonads make mice susceptible to isolation-induced hyperglycemia and increased levels of gluconeogenic glucagon. Additionally, we identify differences in isolation-induced modulation of vlVMH activity.

## Results

### Social isolation induces fasting hyperglycemia in male but not female mice

We used young adult mice (8-10 weeks old) that had been co-housed with same-sex and same-age conspecifics for at least one week after arrival in our facility. We measured blood glucose in mice before (PRE) and after (POST) six hours of fasting at baseline and then after 3 weeks of co-housing or single-housing. In co-housed males we detect significantly lower levels of glucose post-fasting after 3 weeks of co-housing and testing (**Fig. 1a**, Baseline PRE: 150.6 ± 14.23 mg/dl, Baseline POST: 137.0 ± 16.68 mg/dl, 3 weeks of CH PRE: 161.3 ± 21.54 mg/dl, 3 weeks of CH POST: 142.9 ± 22.64 mg/dl, two-way RM ANOVA effect of fasting p = 0.0133, N = 10). After 3 weeks of single-housing males showed increased post-fasting glucose levels (**Fig. 1b**, Baseline PRE: 143 ± 10.93 mg/dl, Baseline POST: 138.8 ± 14.85 mg/dl, 3 weeks SH PRE: 134.8 ± 12.68 mg/dl, 3 weeks SH POST: 162.3 ± 12.20 mg/dl, two-way RM ANOVA effect of fasting p = 0.0039, effect of housing p = 0.0481, and significant interaction p = 0.0100, N = 10).

**Fig. 1:**
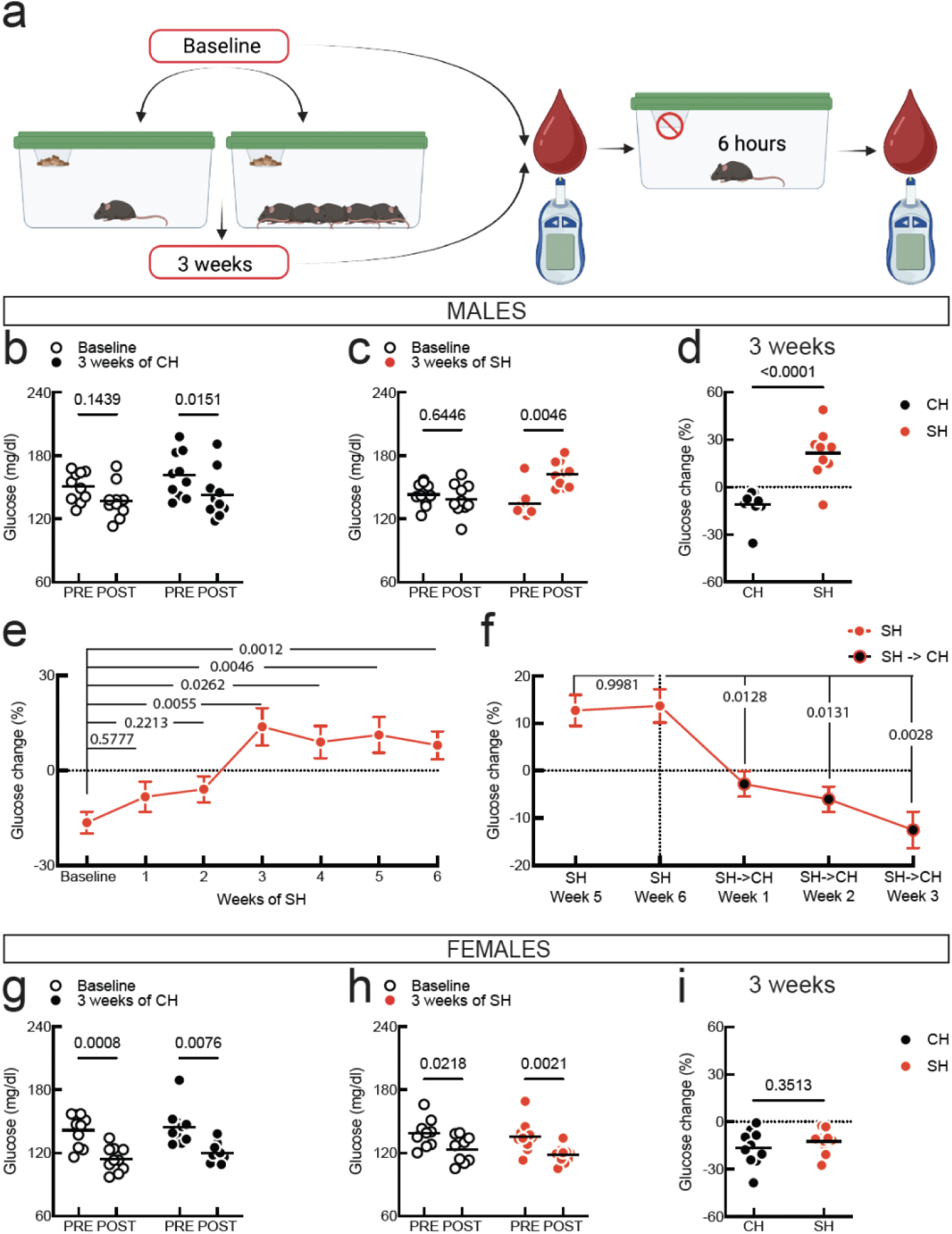
Impact of social isolation on glucose regulation in male and female mice. **a.** Experimental procedure. **b.** Baseline and 3-week data for co-housed (CH) male mice (N = 10). **c.** Baseline and 3-week data for single-housed (SH) male mice (N =10). **d.** Change in blood glucose post-fasting for CH vs SH mice from a and b. **e.** Timeline of glucose changes in male mice after SH onset (N = 14). **f.** Glucose changes in re-grouped mice that had been isolated for 6 weeks (SH→CH). **g.** Baseline and 3-week data for CH female mice (N = 10). **h.** Baseline and 3-week data for SH female mice (N = 10). **i.** Change in blood glucose post-fasting for CH vs SH mice from f and g. For b, c, and g, h, symbols are individual mice, horizontal line is mean, and the pairwise comparisons are Sidak’s tests. For d and i, symbols are individual mice, horizontal line is mean, and pairwise comparison is two-tailed unpaired t-test. For e and f, symbols are mean, errors are SEM, pairwise comparisons are Dunnett’s tests.

We note that at baseline male mice in either group did not show a significant decrease in blood glucose post-fasting, possibly due to insufficient habituation to stress related to blood glucose measurement. Therefore, we focused on glucose dynamics after 3 weeks of either co- or single-housing by calculating the change in glucose post-fasting as a percent of pre-fasting glucose (**Fig. 1c**, CH: −11.15 ± 9.06 %, SH: 21.49 ± 15.56 %, N = 10/housing). Thus, social isolation leads to a remarkable dysregulation of blood glucose, where fasting results in hyperglycemia instead of decreasing glucose levels as in co-housed mice.

To understand the timeline of how fasting hyperglycemia develops in single-housed male mice, we analyzed glucose dynamics every week after isolation onset. We find a progressive alteration of glucose regulation, but only at 3 weeks after the start of single-housing we detect significant and thereafter persistent fasting hyperglycemia (**Fig. 1d**, glucose change at baseline: −16.42 ± 12.91 %, at 1 week of SH: −8.25 ± 18.22 %, at 2 weeks of SH: −5.88 ± 15.32 %, at 3 weeks of SH: 13.86 ± 22.21 %, at 4 weeks of SH: 9.00 ± 18.91 %, at 5 weeks of SH: 11.30 ± 21.26 %, at 6 weeks of SH: 8.031 ± 13.94 %, Mixed-effects model (REML) p = 0.0004, N = 14 mice). To determine if isolation-induced fasting hyperglycemia could be reversed, we re-grouped mice with their old cage mates after 6 weeks of single-housing (SH→CH). We find that one week of co-housing is sufficient to correct fasting hyperglycemia in previously isolated mice (**Fig. 1e**, glucose change after 6 weeks of isolation: 13.64 ± 10.55 %, at one week after SH→CH: −2.81 ± 8.04, two weeks after SH→CH: −6.02 ± 7.98 %, three weeks after SH→CH: −12.49 ± 11.55 %, RM one-way ANOVA p < 0.0001, N =9).

In females, we did not find a significant effect of single-housing on glucose regulation, as blood glucose decreased after fasting at baseline and after 3 weeks of either co-housing or single-housing (**Fig. 1f**, CH - Baseline PRE: 141.5 ± 15.34 mg/dl, Baseline POST: 114.1 ± 11.81 mg/dl, 3 weeks of CH PRE: 144.7 ± 17.67 mg/dl, 3 weeks of CH POST: 119.4 ± 9.07 mg/dl, Two-way RM ANOVA effect of fasting p = 0.0003, N = 10. **Fig. 1g**, SH - Baseline PRE: 138.7 ± 13.95 mg/dl, Baseline POST: 123.3 ± 12.94 mg/dl, 3 weeks of SH PRE: 135.5 ± 14.74 mg/dl, 3 weeks of SH POST: 117.9 ± 7.85 mg/dl, Mixed-effects model REML effect of fasting p = 0.0003, N = 10). The dynamics of glucose regulation in females were therefore unaffected by social isolation (**Fig. 1h**, CH: −16.48 ± 11.17 %, SH: −12.42 ± 7.40, N = 10/housing). These initial findings show that social isolation induces fasting hyperglycemia in males but not females. Furthermore, in males, glucose dysregulation progressively develops over 3 weeks of isolation after which it plateaus, and can be reversed rapidly by re-grouping mice.

### Role of gonadal factors in glucose susceptibility to social isolation

To determine why male mice are susceptible and females resilient to isolation-induced fasting hyperglycemia, we investigated the role of gonads. Male gonadal hormones could enable fasting hyperglycemia in single-housed mice or female gonadal hormones could protect from isolation-induced glucose dysregulation. To test the role of male gonads, we performed orchiectomies or sham surgeries in 8-week old male mice, allowed 2 weeks for recovery and hormonal clearance, and then collected blood glucose at baseline and after 3 weeks of either co-housing or single-housing (**Fig. 2a**). Orchiectomies change baseline glucose levels (**Fig. 2b**, Sham PRE: 168.2 ± 27.41 mg/dl, Sham POST: 147.3 ± 21.55 mg/dl, ORX PRE: 173.1 ± 22.40 mg/dl, ORX POST: 130.4 ± 16.56 mg/dl, two-way RM ANOVA no overall effect of surgery, p = 0.2954, effect of fasting, p < 0.0001, and a significant interaction, p = 0.0137, N =20/group). Orchiectomy prevented isolation-induced hyperglycemia (**Fig. 2c**, glucose change in Sham CH: −11.81 ± 15.43 %, Sham SH: 7.77 ± 14.27 %, ORX CH: −24.89 ± 12.63 %, ORX SH: −19.01 ± 7.23 %, two-way ANOVA effect of housing, p = 0.0036, effect of surgery, p < 0.0001, but no interaction, p = 0.1020, N = 5). Whereas isolation significantly altered glucose change scores in sham mice, it had no effect on glucose dynamics in orchiectomized mice (**Fig. 2c**).

**Fig. 2:**
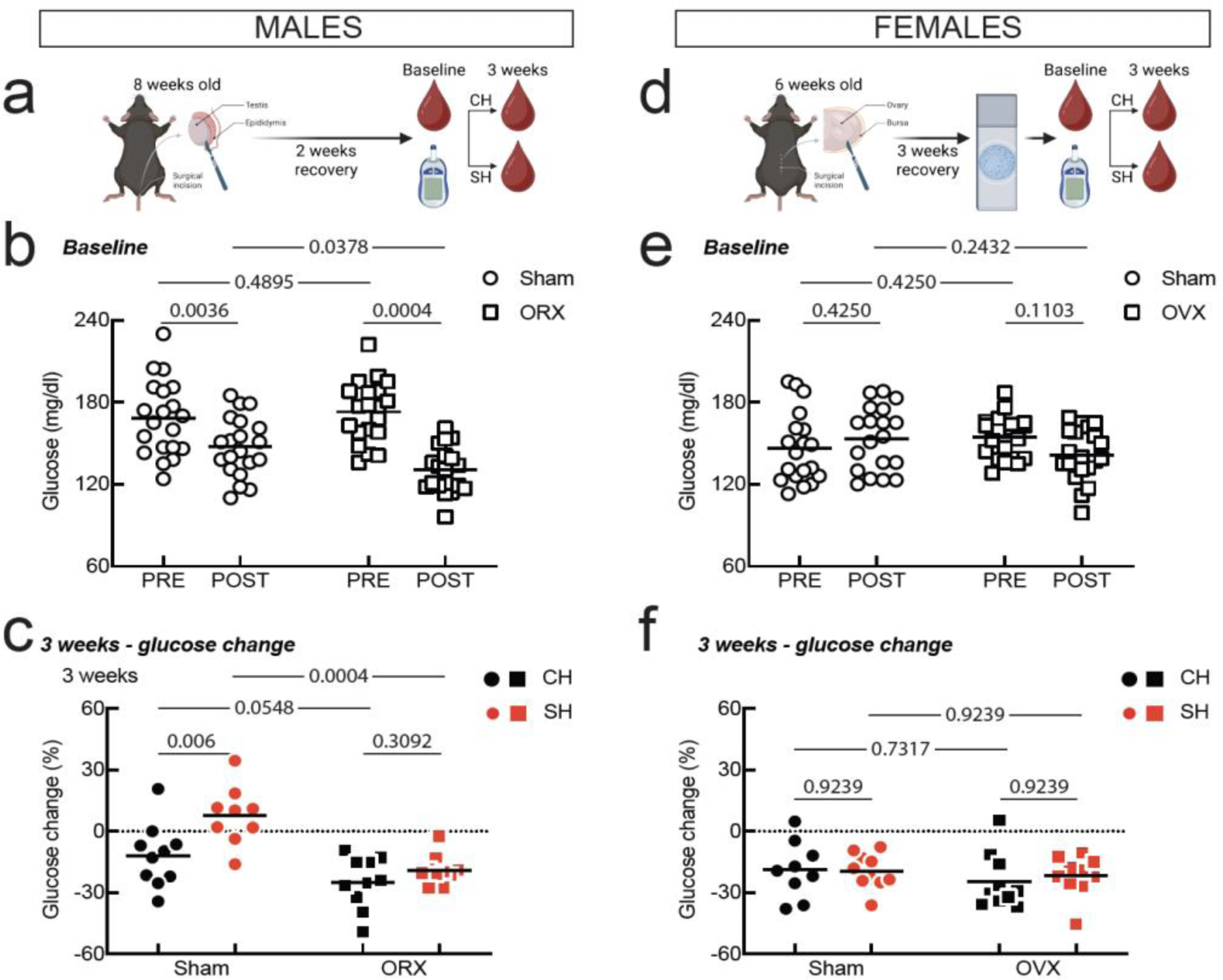
Role of gonads in isolation-induced fasting hyperglycemia. **a.** Orchiectomy (ORX) procedure. **b.** Pre- and post-fasting glucose at baseline for sham and ORX mice (N = 20/group). **c.** Change in blood glucose post-fasting for CH and SH sham and ORX males (N = 10/group). **d**. Ovariectomy (OVX) procedure. **e.** Pre- and post-fasting glucose at baseline for sham and OVX mice (N = 20/group). **f.** Change in blood glucose post-fasting for CH and SH sham and OVX females (N = 10/group). For b., c., e., and f. symbols are individual mice, horizontal line is mean, and the pairwise comparisons are Holm-Sidak’s multiple correction tests.

To test the role of female gonads, we performed ovariectomies and sham surgeries on 6-week-old female mice, we allowed 3 weeks for recovery and hormone clearance, and then confirmed successful ovariectomies with vaginal cytology (**Fig. 2d**). Unlike sham animals that displayed normal cytology patterns covering the different phases of the estrus cycle, ovariectomized females had vaginal smears depleted of most epithelial cells and with few leucocytes (**Fig. S1a**). Following recovery, we collected baseline glucose and then housed animals either in groups of five or single-housed. Both sham controls and ovariectomized females failed to show lower glucose levels post-fasting, most likely due to increased stress compared to the intact females in Fig. 1. Ovariectomy does not affect baseline glucose levels, except for modestly affecting the presumed effect of stress, as revealed by the interaction between surgery and fasting (**Fig. 2e**, Sham PRE: 146.4 ± 25.91 mg/dl, Sham POST: 153.2 ± 23.38 mg/dl, OVX PRE: 154.2 ± 15.56 mg/dl, OVX POST: 141.5 ± 19.17 mg/dl, two-way RM ANOVA no effect of surgery, p = 0.7234, no effect of fasting, p = 0.4665, but a significant interaction, p = 0.0196, N =20/group). Similarly, both sham and ovariectomized mice were resilient to isolation-induced glucose dysregulation (**Fig. 2f**, glucose change in Sham CH: −18.78 ± 13.82 %, Sham SH: −19.47 ± 8.60 %, OVX CH: −24.63 ± 13.50 %, OVX SH: −21.70 ± 9.91 %, two-way ANOVA no effect of housing, p = 0.7651, no effect of surgery, p = 0.2849, and no interaction, p = 0.6302, N = 10/group). These data show that ovarian hormones do not protect females from the effects of social isolation on glucose regulation. Instead, sex differences in isolation-induced fasting hyperglycemia are driven by testicular hormones.

In addition to glucose dynamics, we also monitored isolation-induced changes in body weight as a function of gonadal status. Isolation does not affect body weight in males or females, and gonadectomy increases body weight only in females (**Fig. S2**). This indicates that social isolation does not significantly affect body weight, although it could affect body composition and other metabolic endpoints in addition to glucose regulation.

### Hormonal changes with social isolation

For the rest of the analysis, we focused on males to mechanistically investigate factors that contribute to isolation-induced hyperglycemia and that are sensitive to orchiectomies. We first measured pancreatic hormone differences in cardiac blood collected at the 3-week time point after 6 hours of fasting. Isolation did not change plasma levels of insulin, a major gatekeeper of glucose homeostasis. Although orchiectomies significantly decreased insulin levels, the effect was present in both co-housed and single-housed mice (**Fig 3a**, Sham CH: 0.17 ± 0.11 ng/ml, Sham SH: 0.25 ± 0.14 ng/ml, ORX CH: 0.10 ± 0.04 ng/ml, ORX SH: 0.07 ± 0.07 ng/ml, two-way ANOVA no effect of housing, p = 0.4421, significant effect of surgery, p = 0.0005, but no interaction, p = 0.1374, N = 37).

**Fig. 3:**
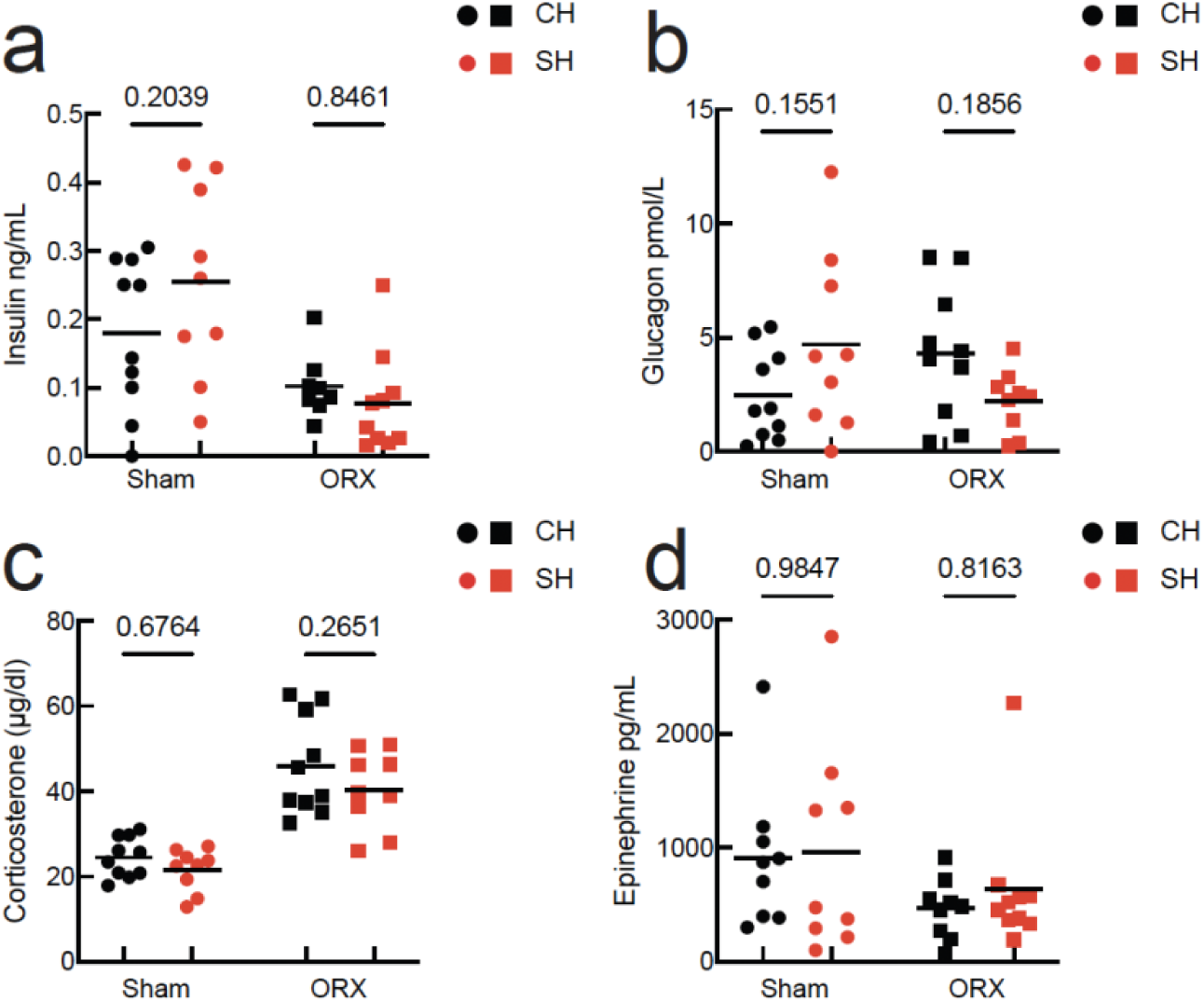
Modulation of glucoregulatory hormones by social isolation. **a.** Post-fasting insulin levels in CH and SH intact and ORX males (N = 10/group). **b.** Glucagon levels. **c**. Corticosterone levels. **d.** Epinephrine levels. Symbols are individual mice, horizontal line is mean, and the pairwise comparisons are Holm-Sidak’s multiple correction tests.

Glucagon, however, was differentially affected by single-housing in sham and orchiectomized mice (**Fig. 3b**, Sham CH: 2.47 ± 1.96 pmol/L, Sham SH: 470 ± 3.93 pmol/L, ORX CH: 4.33 ± 2.89 pmol/L, ORX SH: 2.22 ± 1.36 pmol/L, two-way ANOVA no effect of surgery, p = 0.7242, no effect of housing, p = 0.9464, but a significant interaction, p = 0.0184, N = 38).

Pancreatic α-cells release glucagon in response to stress hormones such as corticosterone and epinephrine, which also have direct hepatic gluconeogenic effects (Barseghian & Levine, 1980; De Marinis et al., 2010; Granner et al., 1984; Saccà et al., 1983). Single-housing enables physiological stress via the activation of the hypothalamo-pituitary-adrenal axis which results in increased release of gluconeogenic corticosterone hormone (Heck et al., 2020), or via sympathetic activation which results in increased release of epinephrin (Cannon, 1915; Mumtaz et al., 2018). However, we did not detect significant differences in corticosterone between co-housed and single-housed intact mice, and orchiectomy equally increased corticosterone levels in both housing conditions (**Fig. 3c**, Sham CH: 24.52 ± 4.64 µg/dl, Sham SH: 21.55 ± 4.92 µg/dl, ORX CH: 45.98 ± 11.51 µg/dl, ORX SH: 40.38 ± 9.07 µg/dl, two-way ANOVA effect of surgery, p < 0.0001, no effect of housing, p = 0.1136, and no interaction, p = 0.6220). Similarly, we did not detect significant differences in plasma epinephrine levels (**Fig. 3d**, stat to be added, Sham CH: 913.8 ± 643.2 pg/ml, Sham SH: 961.4 ± 912.7pg/ml, ORX CH: 465.4 ± 261 pg/ml, ORX SH: 634.4 ± 592.0 pg/ml, two-way ANOVA no effect of surgery, p = 0.0763, no effect of housing, p = 0.6125, and no interaction, p = 0.7762). These data identify glucagon as the major gluconeogenic hormone modulated by the interaction of social isolation and testicular factors.

### The effects of social isolation on neuronal substrates for glucose homeostasis

In addition to circulating hormones, glucoregulatory brain structures can modulate glucagon release via polysynaptic circuits (Faber et al., 2018; Fioramonti et al., 2010; Rosario et al., 2016). The PVN represents a key structure at the intersection of glucoregulatory and social behavior brain networks and connects with the autonomic innervation of the pancreas (Papazoglou et al., 2022; Rosario et al., 2016). Since acute isolation increases the activity and abundance of PVN oxytocin-expressing neurons (Musardo et al., 2022), and these neurons were shown to regulate pancreatic activity (Papazoglou et al., 2022), we investigated if chronic isolation affects the number of oxytocin-expressing neurons in PVN. We did not detect a significant difference between co-housed and single-housed animals in the proportion of PVN oxytocin neurons. (**Figure S3**).

We then investigated if isolation could impact oxytocinergic modulation of vlVMH, a classical glucoreguatory brain structure. Specifically, activation of VMH nNOS1+ neurons induces glucagon release and gluconeogenesis (Faber et al., 2018). Using in situ hybridization RNAscope we find that a substantial proportion of nNOS1+ neurons in vlVMH express the receptor for oxytocin (OTR), indicating that vlVMH-OTR+ neurons could play a role in glucose regulation (**Figure 4a,b**). To determine if social isolation engages vlVMH-OTR+ neurons, we performed in situ hybridization for OTR and the immediate early gene Npas4 in the vlVMH of mice co-housed or single-housed and food deprived for 6 hours. We did not find a significant difference between groups in overall Npas4 or OTR-expressing neurons (**Figure S4**). However, we find that social isolation increases the proportion of OTR+ neurons among activated cells (**Figure 4c,d**). Our histological findings indicate that OTR neurons in vlVMH could play a role in isolation-induced glucose dysregulation.

**Fig. 4:**
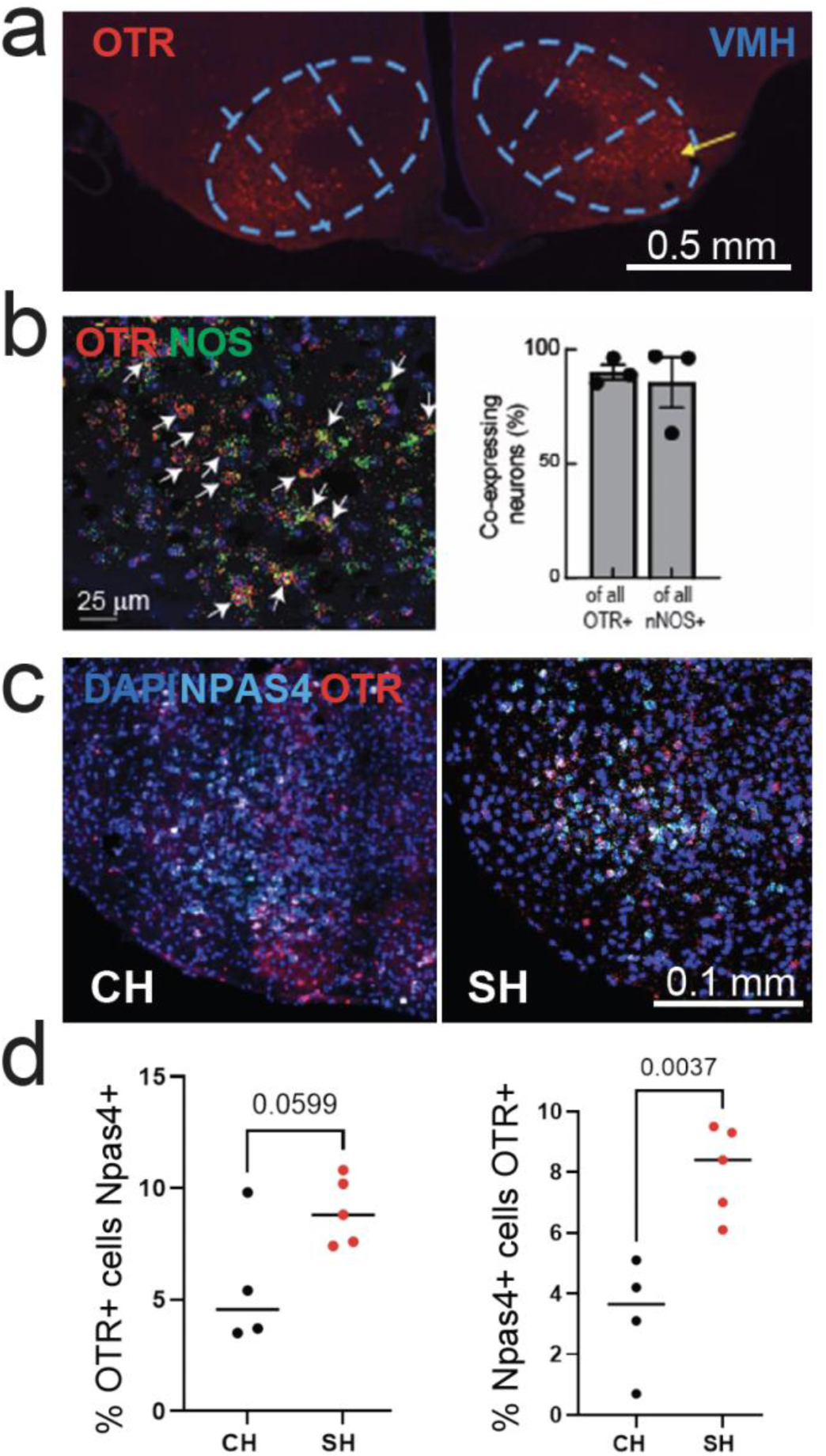
Effects of social isolation on glucoregulatory neurons in vlVMH. **a.** Immunohistochemistry from an OTR-T2A-TdTomato mouse showing OTR expression in vlVMH. **b.** In situ hybridization for OTR and nNOS1 shows significant co-localization (> 80%) in vlVMH. **c**. Example in situ hybridization for OTR and Npas4 in co-housed and single-housed mice. **d.** The effects of isolation on OTR and Npas4 co-expression (N = 4 CH and 5 SH mice, unpaired t-test).

## Discussion

Our work demonstrates that chronic social isolation induces fasting hyperglycemia specifically in intact male mice after 3 weeks, with effects persisting through 6 weeks of isolation. These data extend our previous finding that we were unable to induce hypoglycemia in single housed male mice despite extremely high insulin doses (Patel et al., 2023). Our findings are also in agreement with recent work showing higher fasting glucose in isolated vs co-housed mice (Chen et al., 2025). However, by monitoring both pre- and post-fasting glucose, we demonstrate that fasting hyperglycemia in isolated mice likely results from new glucose production and not from insulin resistance. The temporal dynamics align with previous literature showing chronic but not acute isolation leads to metabolic dysfunction (Sun et al., 2014), though we extend these findings by demonstrating reversibility within one week of regrouping. The differential glucagon response in single vs group housed mice without altered insulin, indicates that engagement of pancreatic α-cells could drive fasting hyperglycemia during chronic isolation. This is further supported by a lack of effect of isolation on epinephrine and corticosterone, although prior work found that chronic isolation impacts the HPA axis (Grippo et al., 2007; Smolensky et al., 2024).

Female mice demonstrate resilience to isolation-induced hyperglycemia, maintaining appropriate fasting hypoglycemia throughout the experimental period. This sex difference parallels human epidemiological data showing stronger associations between social isolation and diabetes risk in men (Lukaschek et al., 2017). Several reasons could explain the sexual dimorphism. First, males preferentially utilize glucose during fasting and in the initial stages of exercise while females rely more on lipid oxidation (Hedrington & Davis, 2015). This is consistent with the observation that activation of vlVMH nNOS glucose-inhibited neurons in low glucose is decreased in females compared to males (Santiago). As a result of this difference in fuel dependence, isolation may exert the same metabolic stress in females which is not detected when using glucose as a primary endpoint. Therefore, future isolation studies should investigate whether isolated females show increased free fatty acid metabolism compared with co-housed females. Another possibility is that female mice have a different ‘threshold of sociability’ and can maintain resilience to the stress of social isolation. However, the former explanation is more plausible given the extensive social isolation literature in female rodents showing that social isolation exerts negative effects on physiology (Grippo et al., 2007; Liu et al., 2025; Normann et al., 2021).

Our findings challenge assumptions about sex hormone roles in metabolic protection. Ovariectomy did not sensitize females to isolation-induced hyperglycemia, while orchiectomy conferred resilience to males, revealing testosterone as permissive of isolation-induced hyperglycemia rather than ovarian hormones as protective. Interestingly, we previously found sex differences in the glucose sensitivity of vlVMH nNOS dependent glucose-inhibited neurons from pre-pubescent male and female mice (Fioramonti et al., 2010; Hirschberg et al., 2020; Santiago et al., 2016b). This further supports ovarian hormone independent sex differences in glucose regulation. On the other hand, estrogen blunts activation of vlVMH glucose-inhibited neurons in low glucose in sexually mature mice and glucose sensitivity varies with the estrus cycle (Santiago et al., 2016a). Together, these observations provide further support for the hypothesis above suggesting that while social isolation causes dysregulation in glucose homeostasis in males, it may instead impact fatty acid oxidation in females and cause metabolic dysfunction.

Importantly, social isolation itself modulates testicular function. Individually housed male mice show significantly increased morning testosterone concentrations and elevated testis weight compared to group-housed controls in both C57BL/6 and BALB/c strains (Sayegh et al., 1990). This isolation-induced testosterone elevation may amplify metabolic vulnerability through multiple mechanisms. The circadian specificity of this effect—occurring only in morning samples—suggests isolation disrupts the normal diurnal regulation of the hypothalamic-pituitary-gonadal axis.

One possibility is that testosterone fundamentally shifts metabolic substrate preference toward glucose utilization and away from lipid oxidation, creating a metabolic state that is vulnerable to perturbation during isolation stress. It is plausible that orchiectomy protects against isolation-induced glucose dysregulation by shifting metabolism toward a female-typical pattern of enhanced lipid oxidation and reduced glucose dependence. This metabolic reprogramming reduces the demand for glucose mobilization during stress, thereby preventing the hyperglycemic responses observed in intact isolated males. Orchiectomized males have been shown to adopt female-typical fat oxidation patterns, which would explain both their lower baseline glucose levels and their resistance to isolation-induced glucose mobilization that we observed (Xu et al., 2018). This substrate shift could represent an adaptive metabolic reprogramming that protects against the glucose dysregulation triggered by social isolation stress in intact males.

The co-localization of oxytocin receptors with vlVMH nNOS-expressing neurons, the majority if not all of which are glucose-inhibited, represents a novel finding potentially linking social peptide systems with glucose sensing. The increased *npas4/otr* co-expression without receptor changes suggests enhanced activation of oxytocin-responsive neurons, possibly compensating for reduced endogenous oxytocin during isolation. Our observation of increased neuronal activation markers without changes in total receptor numbers parallels prior findings (Ross et al., 2019) of altered receptor binding patterns rather than simple expression changes. This suggests isolation induces functional reorganization of hypothalamic circuits—increasing receptor sensitivity and neuronal responsiveness without necessarily changing protein expression levels. Importantly, all brains (single and co-housed) in our study were collected after a six-hour fast, meaning observed differences reflect chronic housing effects on responses to acute stress. The absence of PVN neuropeptide changes suggests metabolic effects operate through functional signaling alterations rather than expression changes.

In summary, stress-induced hyperglycemia is an important evolutionary design which enables endogenous fuel utilization for physical threats. However, modern psychological stress activates these mechanisms without physical activity to utilize mobilized glucose, leading to pathology. The sex-dependent nature of these responses reflects different evolutionary pressures—territorial males requiring rapid glucose mobilization versus communal females prioritizing metabolic efficiency.

## Material and Methods

All procedures were conducted in accordance with institutional animal care guidelines and were approved by the Rutgers Animal Care and Use Committee. C57BL/6N mice were obtained from Taconic Biosciences and delivered weekly to the facility. Animals arrived at 7-8 weeks of age and were allowed a minimum 5-day acclimation period before experimental procedures began. Both male and female mice were used for vivarium experiments, while only males were used for behavior core experiments. Mice were maintained at 22°C ambient temperature with a 12-hour light/dark cycle (lights on at 7:00 AM) and 40-60% relative humidity. Food (standard chow diet, LabDiet 5001) and water were provided ad libitum unless fasting was required for experimental procedures. All experiments were performed during the light phase.

### Blood Glucose Measurements

Two-three days prior to experimental onset, mice underwent habituation procedures including daily handling for approximately 5 minutes per animal with gentle stroking using a gloved hand. This habituation protocol was designed to minimize stress-induced effects on glucose measurements. Initial tail tip blood samples (approximately 1 mm of tissue) were collected during this habituation period to acclimate mice to the tail snip procedure, thereby reducing acute stress responses during subsequent glucose monitoring.

Blood glucose was measured with a glucometer before and following a 6-hour fast at baseline and weekly thereafter for 6 weeks. We chose a moderate fasting interval based on prior literature indicating that at 4-6h time point glucose levels decrease but ketone bodies are stable (Fu et al., 2024). To minimize circadian variation, measurements were collected at standardized times. Pre-fasting measurements were recorded between 10:00-11:30 AM. Post-fasting measurements were recorded between 4:00-5:30 PM. For fasting procedures, mice were gently stroked 2-3 times with a gloved hand to remove food particles from their coat, then transferred individually to clean cages with water but no food. All mice, including group-housed controls, were single-housed during the 6-hour fasting period to standardize conditions and enable individual glucose measurements. Blood glucose was measured from tail blood (5 μL) obtained by massaging the tail from base to tip to reopen the scab from initial tail snip. Measurements were performed using a calibrated Contour EZ glucometer with Contour glucose test strips (blue package). Body weight was recorded after post-fasting glucose measurements to avoid stress-induced effects on glucose levels. Following measurements, group-housed mice were carefully regrouped only with their original cage-mates.

### Social Isolation

For social isolation, 8-10-week-old mice were single-housed in standard polycarbonate cages (28 × 17 × 12 cm) with corncob bedding (1/8 inch, Envigo) and one compressed cotton nesting square (5 × 5 cm, Ancare Corp.) for 6 weeks. Control mice remained group-housed (5 mice per cage). Single-housed mice were handled with cleaned gloves between animals to avoid scent transfer from previous cage-mates. A subset of single-housed mice was regrouped at week 6 with their original cagemates (5 mice per cage) for an additional 3 weeks to assess reversibility of isolation-induced changes. Male mice were used for all social isolation experiments unless otherwise specified.

### Ovariectomies

Bilateral ovariectomy or sham surgery was performed on 6-week-old female mice under isoflurane anesthesia (2-3% for induction, 1.5-2% for maintenance). Mice were positioned in dorsal recumbency, and the surgical area was shaved and disinfected with alternating applications of chlorhexidine and 70% ethanol. A single midline dorsal incision (1 cm) was made caudal to the posterior border of the ribs. The muscle wall was incised, and the ovarian fat pad was exteriorized to expose each ovary. For ovariectomized mice, the ovarian artery and vein were ligated with 5-0 absorbable suture before ovary removal. For sham-operated controls, ovaries were exteriorized and replaced without removal. The muscle layer was closed with 5-0 absorbable suture, and the skin incision was closed with wound clips. Buprenorphine (0.05-0.1 mg/kg, s.c.) was administered for postoperative analgesia immediately following surgery and every 12 hours for 48 hours. Mice recovered for 3 weeks in group housing before experimental procedures. Successful ovariectomy was confirmed by vaginal cytology performed daily for 2-3 consecutive days beginning 3 weeks post-surgery. Vaginal lavage was performed with 10 μL of sterile PBS, and cells were transferred to glass slides, air-dried, and stained with 0.1% crystal violet for 1 minute. Ovariectomy was confirmed by persistent diestrus characterized by predominant leukocytes (>90%) and absence of cornified epithelial cells. Sham-operated controls showed normal 4–5-day estrous cycling. Following confirmation of surgical success, baseline glucose measurements were collected, and mice were then assigned to single or group housing conditions for 6 weeks.

### Orchiectomies

Bilateral orchiectomy was performed on 8-week-old male mice under isoflurane anesthesia (2-3% for induction, 1.5-2% for maintenance, delivered via nose cone). Following anesthetic induction, the scrotal area was shaved and optionally treated with depilatory cream (Nair) for 30 seconds to remove fine hair, followed by cleaning with 70% ethanol. The surgical site was prepared with alternating applications of antiseptic scrub (Betadine or chlorhexidine) and 70% ethanol. A midline scrotal incision (0.5 cm) was made to exteriorize each testis with associated epididymis and vas deferens. The spermatic cord and associated blood vessels were ligated with 4-0 absorbable suture or cauterized per institutional policy, and the testis and epididymis were excised. The procedure was repeated for the contralateral testis. The incision was closed with wound clips or 4-0 absorbable suture. Sham-operated controls underwent identical procedures without testis removal. Buprenorphine was administered preoperatively. Mice were monitored for at least 3 days post-operatively with blue post-operative cage cards. Animals recovered in warmed cages with accessible food and water. Mice recovered for 2 weeks before baseline glucose measurements and subsequent housing manipulations. Successful orchiectomy was confirmed by seminal vesicle atrophy at tissue collection.

### Tissue collection and processing

Tissue collection was performed after either 3 or 6 weeks in experimental housing conditions. Collection occurred at least 2 days after the final glucose measurement to avoid acute stress effects. All mice were fasted for 6 hours prior to tissue collection to standardize metabolic state. All tissue collections occurred between 3:00 PM and 5:00 PM to minimize circadian variation.

Following deep anesthesia with ketamine/xylazine (0.2 mL/mouse of standard mixture, i.p.), terminal blood collection was performed via cardiac puncture. Using a 3 mL syringe affixed with a 25-gauge needle, blood was collected through exsanguination. The needle was inserted at a 30-degree angle just to the left of the xiphoid process, directed toward the left shoulder. Blood (0.5 – 1ml) was withdrawn slowly to prevent cardiac collapse and immediately dispensed into serum separator tubes (MiniCollect ® tube 0.8 ml, Greiner 450472. Blood for serum analysis was allowed to clot at room temperature for 2 hours, then centrifuged at 1,000 x g for 20 minutes at 4°C. Serum was separated, aliquoted into labeled yellow-cap microtubes, and stored at −80°C until analysis.

In a separate cohort, mice were transcardially perfused with 20 mL ice-cold phosphate-buffered saline (PBS, pH 7.4) followed by 20 mL of 4% paraformaldehyde in PBS. Brains were removed and post-fixed for 4 days at 4°C, then cryoprotected sequentially in sucrose solution. Brains were embedded in OCT (Tissue-Tek) and stored at −80°C until sectioning.

### ELISA hormonal measurements

Serum hormone levels were measured using commercially available ELISA kits according to the manufacturers’ instructions. Serum was collected in serum-separating tubes, allowed to clot for 20 minutes at room temperature, centrifuged at 1,000 × g for 20 minutes at 4°C, and aliquots were stored at −80°C until analysis. Samples were thawed on ice immediately before use and were loaded undiluted for all assays. All standards and samples were run in duplicate. Plates were read at 450 nm with background subtraction at 570 nm. Hormone concentrations were calculated from four-parameter logistic (4-PL) standard curves generated in GraphPad Prism.

Glucagon was quantified using the Ultrasensitive Glucagon ELISA (ALPCO, #48-GLUHUU-E01). Insulin was measured using the Ultra Sensitive Mouse Insulin ELISA Kit (Crystal Chem, #90080). Norepinephrine (noradrenaline) was measured with the High Sensitive ELISA (Eagle Biosciences, #50-305-340). Corticosterone levels were quantified with the Corticosterone Competitive ELISA Kit (Invitrogen, #EIACORT). Epinephrine was measured using the Epinephrine ELISA Kit (Colorimetric) (Novus Biologicals, #NBP2-62867). Arginine-vasopressin (AVP) was measured using the Arg8-Vasopressin Competitive ELISA Kit (Invitrogen, #EIAAVP). Thyroxine (T4) levels were quantified using the Thyroxine (T4) Competitive ELISA Kit (Invitrogen, #EIAT4C).

### Immunohistochemistry

Cryoprotected brains were sectioned at 18-25 μm thickness using a cryostat (Leica CM3050S) maintained at −20°C. Sections were collected on positively charged slides (Superfrost Plus, Fisher Scientific) and stored at −80°C until processing. Sections were thawed at room temperature for 20 minutes and washed in PBS. Tissue was permeabilized and blocked in 10% bovine serum albumin (BSA) with 0.1% Triton X-100 in PBS for 90 minutes at room temperature. Primary antibodies diluted in 5% BSA with 0.01% Triton X-100 were applied for 24-48 hours at 4°C. The following antibodies were used: mouse anti-oxytocin (1:100, gift from H. Gainer, NIH); mouse anti-vasopressin (1:100, gift from H. Gainer, NIH); rabbit anti-c-Fos (1:500, Cell Signaling Technology, #2250); rabbit anti-tdTomato (1:500, Takara Bio, #632496); rabbit anti-nNOS (1:200, Cell Signaling Technology, #4231). Following primary antibody incubation, sections were washed 3 × 10 minutes in PBS and incubated with appropriate secondary antibodies (Jackson ImmunoResearch) at 1:500-1:1000 dilution for 1.5-4 hours at room temperature. The following secondary antibodies were used: goat anti-rabbit Cy3; goat anti-mouse Cy2; donkey anti-rabbit Alexa Fluor 488. Nuclei were counterstained with DAPI (1:10,000) for 5 minutes. Slides were mounted with Fluoromount-G (Invitrogen).

### RNAscope in situ hybridization

RNAscope Multiplex Fluorescent V2 kit (Advanced Cell Diagnostics, #323100) was used according to manufacturer’s instructions with modifications for fresh-frozen tissue. Fresh-frozen sections (18 μm) were fixed in 4% PFA for 15 minutes at 4°C, dehydrated through graded ethanol series (50%, 70%, 100%, 100%; 5 minutes each), and treated with Protease IV for 30 minutes at room temperature. Target probes were obtained for oxytocin receptor (Oxtr, #412171), Vasopressin receptor 1A (Avpr1a, #404841), and neuronal PAS domain protein 4 (Npas4, #423431), and neuronal nitric oxide synthase (Nos1/nNOS, #437651)

Custom-designed probe sets were applied targeting mRNAs with the following channel assignments: Channel 3 for Avpr1a/nNOS (green fluorescence), Channel 1 for Oxtr (red fluorescence), and Channel 2 for Npas4 (cyan fluorescence). Probes were hybridized for 2 hours at 40°C in the HybEZ oven, followed by signal amplification using the AMP 1-3 reagents according to the manufacturer’s protocol. Fluorophores (Opal 520, 570, and 690) were applied at 1:1500 dilution. Sections were counterstained with DAPI for nuclear visualization.

Quality control was performed using positive control probes (Ppib-#313911, Polr2a-#312471, Ubc-#310771) and negative control probe (dapB-#310043) on adjacent sections. RNA quality was considered acceptable when positive control probes showed robust punctate staining and negative control showed minimal background. RNAscope probe signals appeared as punctate dots representing individual mRNA transcripts.

### Image Acquisition

Images were acquired using two systems: Nikon A1R+ HD Confocal Microscope with 405, 488, 567, and 637 nm laser lines. Leica TCS SP8 STED 3× Super-Resolution Microscope for high-resolution detection of transcript signals. Imaging parameters for the ventrolateral ventromedial hypothalamus (vlVMH) included: Objective magnification: 40×, XY resolution: 1024 × 1024 pixels, Z-stack acquisition: 2.57 μm step size with 2 steps above and 2 below the focal plane (5-plane stack from −0.15 μm to 10.13 μm depth), LED_405 excitation for DAPI nuclear staining. All imaging was performed at the Rutgers New Jersey Medical School Imaging Core. For quantification, 2-3 sections per animal were imaged, with 2 fields per section. Acquisition parameters were held constant throughout to ensure signal comparability across experimental groups.

### Quantitative Analysis

Images were analyzed using QuPath software (version 0.6.0), an open-source digital pathology platform for automated quantification of spatially resolved gene expression. For immunofluorescence quantification, cells were identified by DAPI nuclear staining using the cell detection algorithm. Oxytocin and vasopressin neurons were identified by cytoplasmic immunoreactivity. c-Fos-positive cells were identified by nuclear immunoreactivity with intensity >2 standard deviations above background. Colocalization was determined when signals overlapped within the same cell boundary. For RNAscope analysis, DAPI staining was used for cell detection with automated cell segmentation. Transcript puncta were detected using the subcellular detection feature in QuPath. Background threshold was established using negative control sections. Cells were considered positive for a given transcript when containing ≥5 puncta, based on negative control analysis showing <5 puncta per cell in dapB-treated tissue. Co-expression was defined as overlapping fluorescent signals for multiple markers within the boundary of a single cell, identified using the classifier function. Quantitative outputs included: total region of interest (ROI) area, marker-specific cell counts and percentages, number and percentage of co-expressing cells (e.g., NPAS4+/OTR+, OTR+/NPAS4+). Data were exported from QuPath to Microsoft Excel for automatic calculation of cell counts and co-expression percentages. Poor-quality images or those with significant artifacts were excluded from analysis. All images were aligned to anatomical landmarks for the ventrolateral ventromedial hypothalamus.

## Acknowledgements

We would like to thank Jiadong Li for helping with orchiectomies, Samantha Brewer, Stephanie Garcia, James Muldowney and Vasisht Yegneshwaran for helping with tissue collection. We thank the Rutgers Imaging Core for help with histological experiments. We thank Ali Yasrebi, Dr. Gwyndolin Vail, Dr. Pallabi Sarkar for technical assistance, Dr. Troy Roepke for feedback throughout the project, and Dr. Andrew Thomas for support and guidance.

## Authors Contribution

HHL, VHR, and IC co-designed and funded the study, analyzed the data and wrote the manuscript. HHL, RO, PDM and PB performed most of the experiments in the study. RO and PDM helped editing the manuscript.

## Supplementary figures

**Fig. S1:**
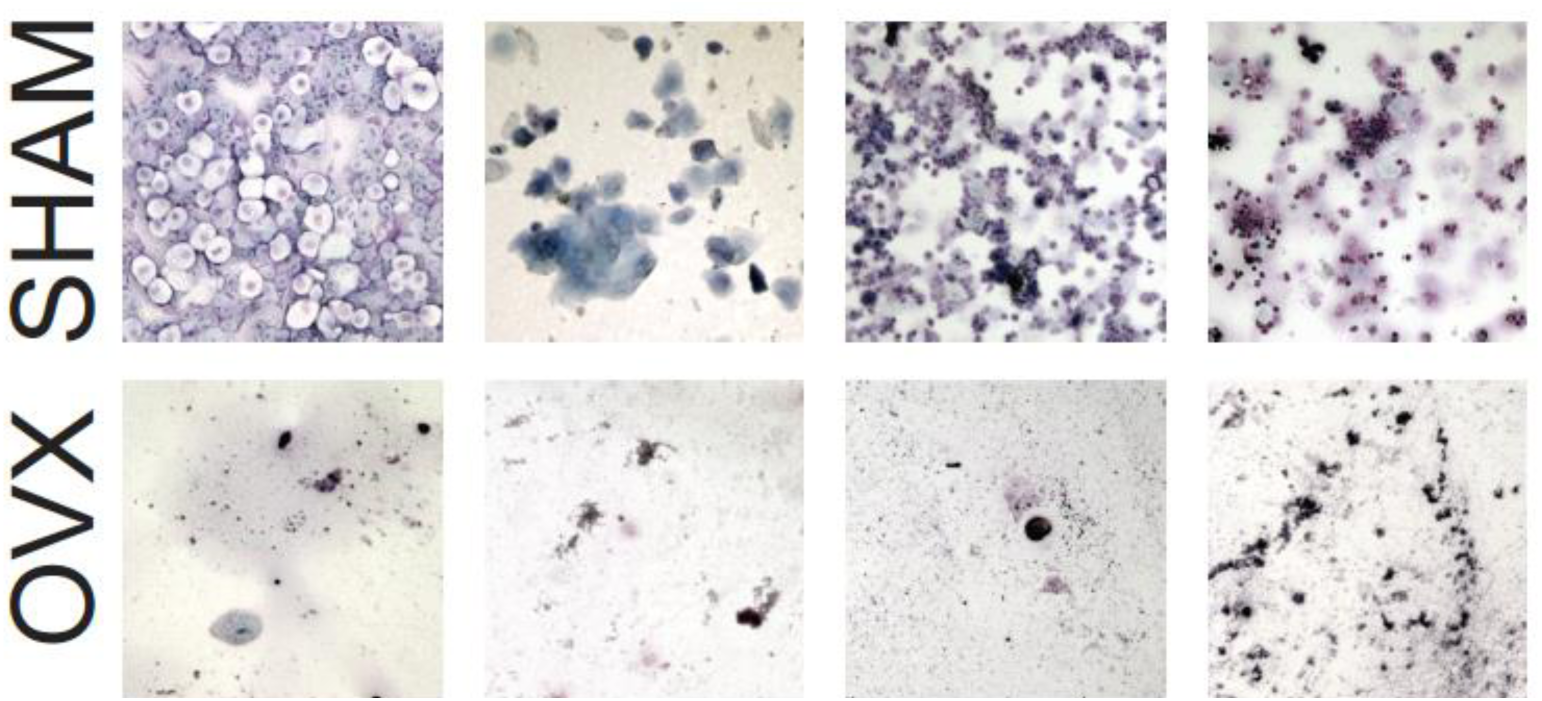
Example of vaginal cytology confirming successful ovariectomy in females. **Top,** cytology showing different phases of the estrus cycle in sham operated animals. **Bottom**, cytology from OVX females, showing minimal cells.

**Fig. S2:**
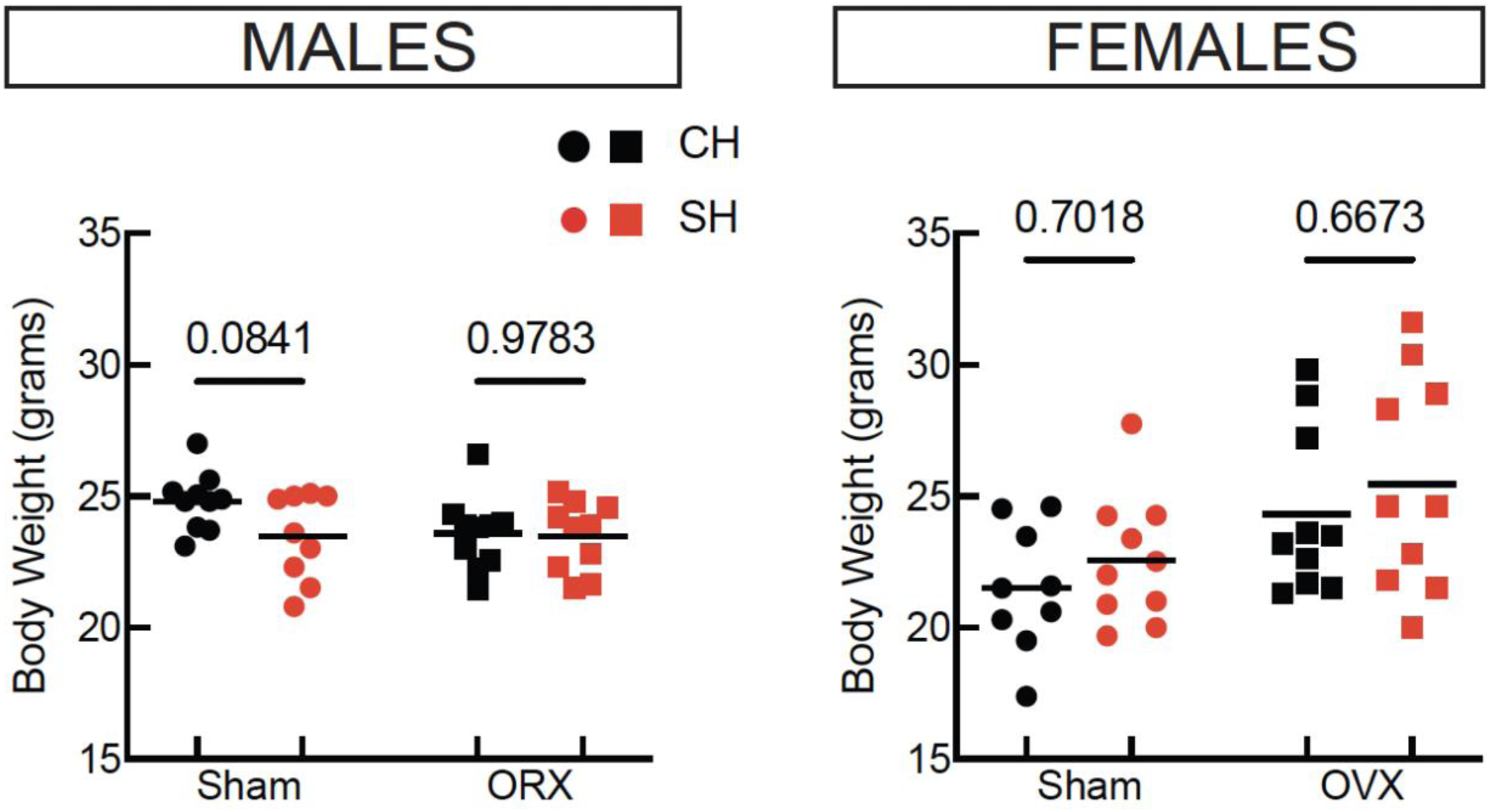
The effects of isolation on body weight in intact and gonadectomized mice. **Left,** no effect of housing, surgery, or their interaction in male mice (two-way ANOVA, N = 19 sham, 20 ORX). In Sidak’s corrected pairwise comparisons there was a trend for lower body weight in isolated compared to co-housed sham animals. **Right**, isolation does not affect body weight in females, but OVX surgery leads to increased body weight (p = 0.007, two-way ANOVA, N = 19 sham, 20 OVX).

**Fig. S3:**
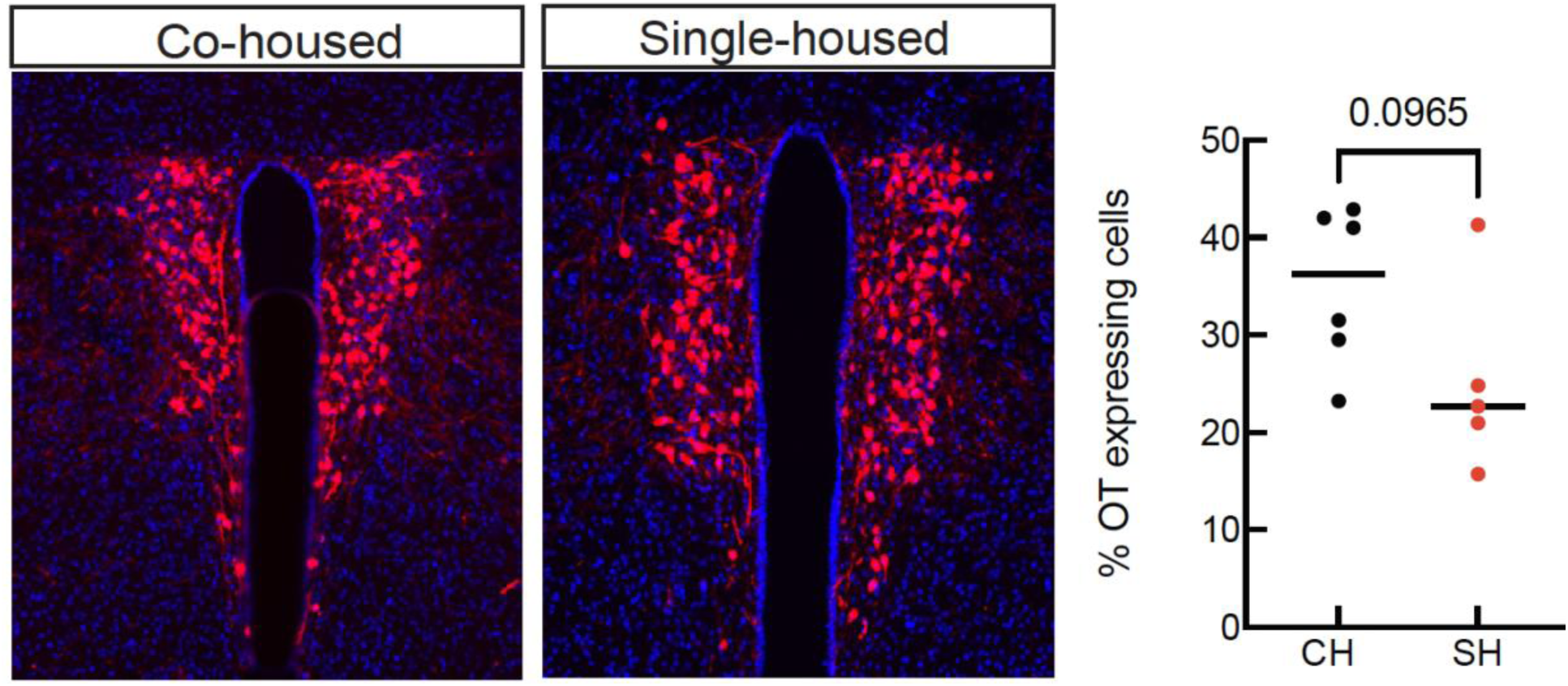
Isolation does not significantly change the proportion of oxytocin neurons in PVN. **Left,** oxytocin neurons (in red) in PVN. Right, quantification of oxytocin neurons in PVN (unpaired t-test, N = 6 co-housed and 5 single-housed male mice).

**Fig. S4:**
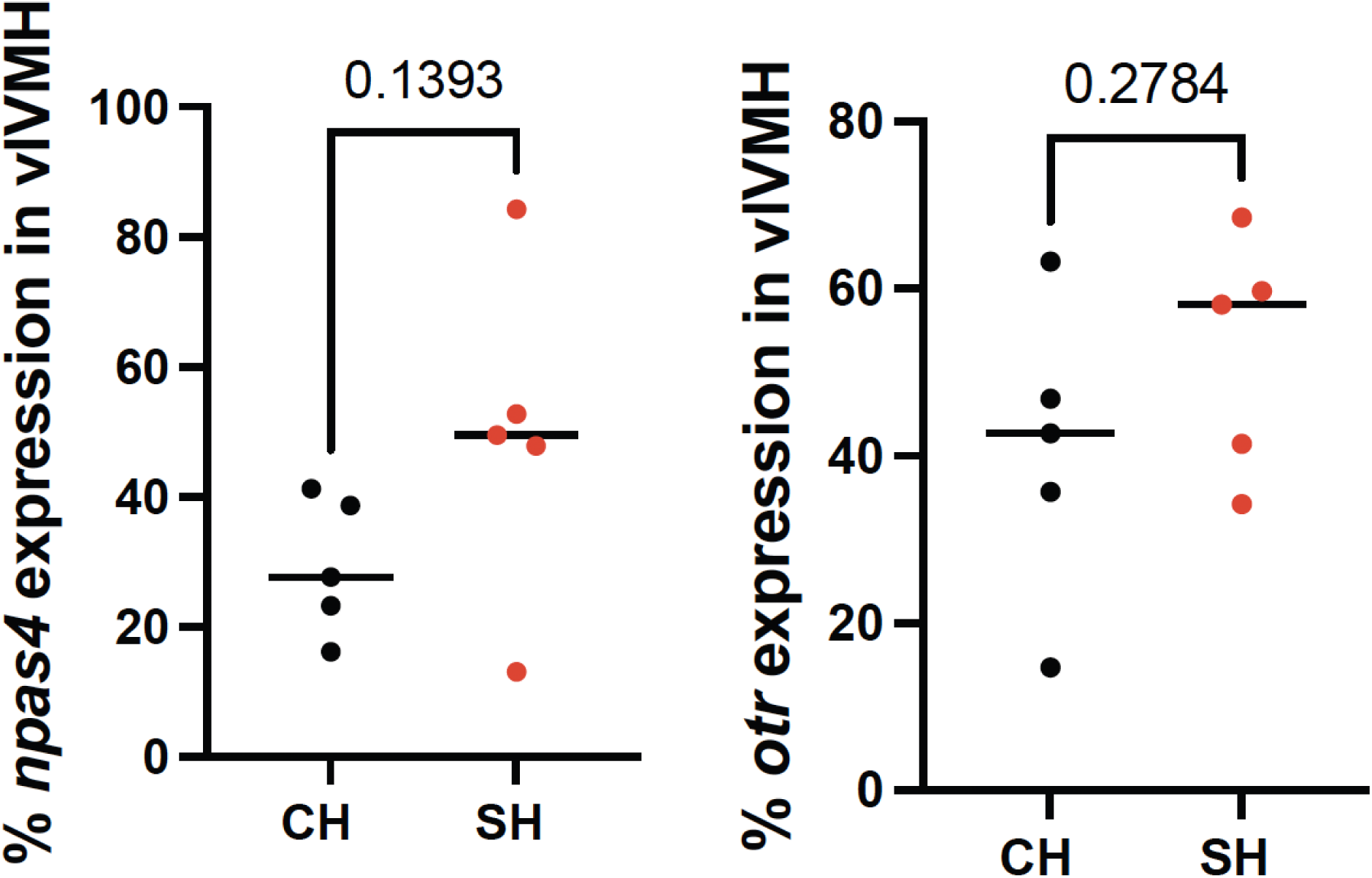
Isolation does not significantly change Npas4 and OTR expression in VMH. Quantification from N = 5 co-housed and 5 single-housed male mice. Unpaired t-test.

